# Effects of *Lactobacillus plantarum* 15-1 and fructooligosaccharide on the response of broilers to pathogenic *Escherichia coli* O_78_ challenge

**DOI:** 10.1101/533935

**Authors:** Sujuan Ding, Yongwei Wang, Wenxin Yan, Aike Li, Hongmei Jiang, Jun Fang

## Abstract

One-day-old broilers were randomly allocated to 5 treatments: basal diet challenged by saline (negative control, n-control); basal diet and challenged by *E.coil* O_78_ (positive control, p-control); supplementation with *L. plantarum* 15-1 at 1×10^8^ CFU/kg challenged with *E.coil* O_78_ (LP); supplementation with FOS at 5 g/kg challenged with *E.coil* O_78_ (FOS); supplementation with *L. plantarum* 15-1 and FOS challenged with *E.coil* O_78_ (LP+FOS). *L. plantarum* 15-1 or FOS had a lowered effect (*P*<0.05) on crypt depth on d 14 compared with two controls, and *L. plantarum* 15-1, FOS and *L. plantarum* 15-1+FOS also reduced relative to p-control on d 21. *L. plantarum* 15-1 reduced the level of diamine oxidase (DAO) at d 14 and 21 compared with p-control (*P*<0.05), the broilers with *L. plantarum* 15-1 and FOS increased the concentration of IgA and IgG relative to two control, and decreased diamine oxidase (DAO) compared with p-control (*P* 0.05). *L. plantarum* 15-1 increased the concentration of acetic acid and total short-chain fatty acid (SCFA) in comparison with p-control at d 14 (*P*<0.05), FOS improved the level of valeric acid and total SCFA relative to p-control at d 21 (*P*<0.001), the broilers fed *L. plantarum* 15-1 and FOS were increased the level of butyric acid at d 14 (*P*<0.05). FOS enhanced bursal index of broilers at d 21 (*P*<0.05). *L. plantarum* 15-1 and FOS did no effect on the growth performance. In conclusion, FOS can promote average daily gain, *L. plantarum* 15-1 and FOS can improve intestinal morphology, and increase the concentration of SCFA in cecal contents in broilers challenged with *E.coil* O_78_. These results suggest that *L. plantarum* 15-1 and FOS have effective mitigation to *E. coil* O_78_ via lowing reducing the intestinal injury and enhancing the immune responses.

## Introduction

*Escherichia coli* (*E. coli*)-induced diarrhoea has become a global public health problem, both in developed and developing countries. At present, the prevention and treatment of the disease is mainly based on drugs and vaccines. In addition, dietary intervention has also become an important means [1]. *E. coli* can produce the enterotoxin that causes colibacillosis, which brings untold damage for the poultry production [2, 3]. *E. coli* whose serotypes are O1, O2, O78, O15 and O55 had been found to be related to colibacillosis in chickens [4, 5], which may undermine immune function to predispose host animals to the colonization of the pathogens, placing the threat to health and food safety. Although antibiotic therapy is effective for colibacillosis, there are increasing restriction and ban to limit the use of antibiotic to poultry. Therefore, prebiotic and probiotic as candidates to replace antibiotics are available, they are more secure to prevent and control colibacillosis, thus protecting livestock species. Short-chain fatty acids produced by intestinal microbiota are one of the important determinants of the interaction between intestinal microorganisms and pathogenic bacteria [6]. A study has shown that the diet supplied with lactulose improved body weight gain and feed conversion efficiency of 21-day-old broilers, but had no effect on growth performance of 42-day-old broilers. In addition, lactulose treatment increased the colonies of Lactobacillus in cecum, and increased the levels of acetic acid, propionic acid, butyric acid and total SCFA in cecum contents of 7-day-old and 42-day-old broilers [7].

Probiotics were defined as live microbial feed supplements, which have positive effect on the host animal by improving intestinal microecology [8]. The probiotic keep a healthy intestinal microflora and stimulate the immune response of the host animal decreasing pathogenic microbiota of the gut [9]. An increasing number of well-characterized probiotic strains have been chosen to prevent pathogen bacteria and thus maintaining a healthy avian intestinal microbiota. Especially numerous researches on feeding *Lactobacillus spp*. to broiler have been increasing to assess their effect on immune function, performance and reduction of pathogen shedding. Experiment in vitro [10] found that lactobacilli inhibited the TLR4 inflammatory signaling induced by enterotoxigenic *Escherichia coli* via modulation of the inflammation and involvement of TLR2 in human intestinal Caco-2/TC7 cells and intestinal explants. *Lactobacillus Plantarum* inhibited the growth of *Escherichia coli* O157:H7 in vitro [11]. Moreover, *Lactobacillus Plantarum* possesses the ability to improve the growth performance, reducing *Enterobacteriaceae*, increasing the population of *Lactobacillus*, the villi height of the small intestine and the concentration of volatile fatty acids in faeces of the broiler [12].

Prebiotics are a non-digestible food or feed ingredients, which positively affect the host by selectively stimulating the growth and activity of one or a limited number of bacteria in the colon [13]. Common prebiotics includes fructooligosaccharide, inulin, galactooligosaccharides (GOS), transgalacto-oligosaccharides (TOS), and lactulose. Intake of prebiotics can regulate the intestinal microbiota by increasing the number of specific probiotic bacteria such as *Lactobacillus* and *Bifidobacterium* [14] or competing with pathogens bacteria for attachment site thus reducing the pathogenic bacteria in the intestinal tract [15]. G.-B. Kim et al.[16] investigated the effect of FOS on growth performance and immune response in broiler chickens. The results showed that adding 0.25% FOS to diets was comparable to avilamycin, reducing the population of *E. coli* and increasing the population of lactobacilli. This study aimed to reduce the negative effect on the intestinal morphology and the decline of immune response induced by *E. coli* O_78_ after administrating with *L. plantarum* 15-1 and FOS.

## Materials and Methods

### Broilers, Diets, and Experimental Design

All animal procedures were approved by the Animal Ethics Committee Guidelines of Academy of State Administration of Grain, Beijing, China (20170052), and performed according to the guidelines recommended in the Guide for the Laboratory Animal Ethical Commission of State Administration of Grain. 150 one-day-old male Arbor Acres (AA), with the average body weight of 46.38 ± 0.13 g, were used in this experiment. The broilers were obtained from a commercial hatchery (Huadu Broiler Farms, Beijing, China). The broilers were randomly allocated to 5 experimental groups (5 broilers per treatment across six replicate pens). The broilers challenged was kept in a separate room, while n-control with other treatments were placed in another room inoculated with the sterilized saline solution as the unchallenged group. The broilers were kept in cages with wire mesh floor, breeding density is 550 cm^2^/broilers. Room temperature was adjusted to 32±2°C in the first week and gradually dropped to 24°C by the end of the third week.

The diets were fed to the four groups, n-control has a same dietary with p-control, namely: 1) n-control and p-control (without any dietary additives); 2) *L. plantarum* 15-1 treatment (1 × 10^8^ CFU/kg of feed); 3) FOS treatment (5 g/kg of feed); 4) *L. plantarum* 15-1 and FOS treatment (1 × 10^8^ CFU/kg and 5 g/kg of feed). *L. plantarum* 15-1 was added to the basal diet as powder, and the final colony count reached 1 × 10^8^ CFU/kg in per kilogram of diet. The components of the basal diet are shown in Table 1, the basal diet based on Chinese feeding standard of chicken [17] was antibiotic-free and meet the nutritional requirements for starter 1-21 days.

**Table 1.** Composition of basal diet

| Item | Content |
| --- | --- |
| Ingredient (%) |  |
| Corn | 55.75 |
| Soybean meal | 36.75 |
| Soybean oil | 2.96 |
| Calcium phosphate | 1.86 |
| Limestone | 1.2 |
| Sodium chloride | 0.35 |
| Lysine | 0.309 |
| Solid methionine | 0.287 |
| Threonine | 0.05 |
| Choline chloride (50%) | 0.26 |
| Minerals premix<sup>1</sup> | 0.2 |
| Vitamin premix<sup>2</sup> | 0.02 |
| Total | 100 |
| Calculated nutrient composition |  |
| ME, kcal/kg | 3200 |
| Crude protein % | 22.16 |
| Ca % | 1.07 |
| Available P % | 0.68 |
| Lys % | 1.32 |
| Methionine | 0.48 |

<sup>1</sup>Vitamin premix provided the following per kilogram of diet; vitamin A, 9,500 IU; vitamin D<sub>3</sub>, 62.50 µg; vitamin K<sub>3</sub>, 2.65 mg; vitamin B<sub>12</sub>, 0.025 mg; vitamin B<sub>2</sub>, 6 mg; vitamin E, 30 IU; biotin, 0.0325 mg; folic acid, 1.25 mg; pantothenic acid, 12 mg; nicotinic acid, 50 mg. <sup>2</sup>The mineral premix provided the following per kg of diet: Cu, 8 mg; Zn, 75 mg; Fe, 80 mg; Mn, 100 mg; Se, 0.15 mg; I, 0.35 mg

### Bacterial Preparation and Oral Challenge

The strain of *L. plantarum* 15-1 used in this study was offered by Academy of State Administration of Grain, Beijing, China. The *L. plantarum* 15-1 added to the basal diet was freeze-dried powder, and the final concentration of group is 1×10^8^ CFU/kg. The number of colonies in freeze-dried powder is determined by calculating the number of colonies on the plate. Briefly, 10g freeze-dried powder was added into 90 ml sterile water, the mixture was diluted 1:10 after mixing evenly, then plated 100µl of dilutions evenly onto the MRS Agar (De Man, Rogosa, Sharpe). The plates were incubated at 37°C for 10-12 h, colonies were counted on the plate. A gram of CFU per g was calculated according to the dilution multiple of the sample.

The *E. coli* O_78_ was obtained from College of Animal Science and Technology, China Agricultural University, Beijing, China. The serotype of *E. coli* O_78_ was analyzed by China Institute of Veterinary Drug Control (Beijing, China). The strain was aerobically cultured in Luria Bertain broth (LB) for 18-24 h at 37°C with shaking 160 rpm. The gradient dilution medium (1:100) was plated evenly on LB solid media with sterile plates, and the whole process operated in a sterile environment. The plates were incubated at between 35 and 37°C for 18 to 24 h. The colonies of *E. coli* O_78_ were counted [18]. The concentration of overnight culture was 3×10^8^ CFU/ml. Each chicken was oral with 1×10^8^ CFU *E. coli* O_78_ using a sterile syringe in the back of the oral cavity at 7-d-old broilers after making 1:3 dilution.

### Growth performance

Broiler chickens were fasted for 12 hours prior to sacrifice on days 14 and 21, and the body weight and feed intake of each broiler were recorded on a cage-by-cage basis. Average daily gain (ADG) and average daily feed intake (ADFI) during 1-14, 14-21 and 10-21, and bursal index at 14 and 21 days were calculated. Mortality was calculated on a full-time basis.

### Sample Collection

At the 14^th^ and 21^st^ day, blood was taken from wing veins after 7-8 h of fasting. Each treatment randomly was chosen six broilers from each cage to slaughter by jugular bleeding. Thereafter, bursa of fabricius drew above the back of the cloaca were weight to calculate. The jejunum was cutting off 1cm in the middle piece to fix with formaldehyde solution for morphology. The caecal content was collected aseptically with sterile plastic tubes, then quickly frozen in liquid nitrogen and stored at −80°C until use.

### Jejunal Morphology

Each jejunum tissue was softly rinsed with 0.9% NaCl; the clean samples were fixated in 10% buffered formalin. To be fixated for some time, the samples were buried in paraffin, cut into slices of 2-5μm and mounted slides to stain with hematoxylin and eosin [19]. Complete intestinal villi were selected to measure its villi height and crypt depth. Villus height was measured from the villus basal lamina to the villus apex and crypt depth is between the base (which is by villi height end) to the crypt: villus transition zone [20].

### IgA, IgG and Diamine Oxidase (DAO)

The blood was collected from wing vein for the quantification of IgA, IgG, and DAO. After serum separation and then centrifuged at 10000 g for 4 min and stored at −20°C. The serum concentration of IgA, IgG and DAO were measured by enzyme-linked immunosorbent assay (ELISA) using the standard Chicken kit (Nanjing Jiancheng Institute of Biological Engineering, Nanjing, China). The specific operating procedures are provided by the manufacturer’s instructions. Briefly, Standard substance and samples were added into wells with dilution of 100 µl, the plate cultured in 37°C for 1 h. Next, the plate was washed three times after drying the well. Biotin-antibody was added into per well, then cultured for 60 min at 37 °C, following the plate was washed 3 times, anti-chicken HRPO was added into wells and cultured for 30 min at 37°C. 3,3,5,5 - tetramethylbenzidine substrate solution was added into every well and cultured for no more than 30 min at 37°C. Their concentrations were measured spectrophotometrically at 450 nm. The regression equation of the standard curve is calculated from the concentration of the standard and the OD value, the OD value of the sample is substituted into the equation to calculate the sample concentration. All the measurements of each variable were performed in the same assay environment to avoid inter-assay variation.

### Cecal short-chain fatty acid

The method of analyzing SCFA is according to Schäfer K [21] with some modifications. Frozen cecal digesta were thawed at 4°C until to around 4°C, then diluted 5-fold with double-distilled water in sterile screw-cap tubes, homogenized, centrifuged (Eppendorf/ Centrifuge, 5810R) at 10000 rpm for 10 min. 2-ethyl butyric acid was used as an internal standard (17.01mmol/L). The sample was determined the concentrations of acetate, propionate, butyrate, valerate, isovalerate, and isobutyrate by gas chromatography (Agilent, 7890B) with a flame ionization detection (FID) and fatted with DB-FF column (30m×0.25mm, diameter = 0.5μm, Agilent Technologies, USA). Nitrogen was supplied as the carrier gas at a flow rate of 40 mL/min. The initial oven temperature was 80°C, maintained for 0.5 min, raised to 130°C at 5°C/min and kept for 2min, then increased to 240°C at 20°C /min, at last held for 1 min. The temperature of the FID and the injection port was 270 and 200 °C, respectively. The flow rates of hydrogen and air as fuel gas and oxidant gas were 40 and 450 ml/min, respectively. 1μL of the sample was injected into GC system for analysis, and the detection time of one sample is 19 min. At last, the concentration of SCFA is the system data multiplied by the dilution factor.

### Statistical Analysis

All statistics were performed using SPSS for Windows. Data were analyzed by multivariate one-way ANOVA for the following main effects of broilers (n=6): broilers average daily gain (ADG), average daily feed intake (ADFI), bursal index, Jejunum villus height, crypt depth, immune responses and the concentrations of SCFA were statistically analyzed with independent Student’s t-test. Differences in the effects of *L. plantarum* 15-1 and FOS were determined with a single degree of freedom contrast statements comparing broiler challenged with *E. coli* O_78_ (p-control) and unchallenged group (n-control) from 0 to 21 d. Differences were considered significant at *P*<0.05.

## RESULT

### Growth Performance and the Survival

The growth performance of broilers showed in Table 2, *E. coli* O_78_ lowered the average weight on the 21^st^ day, decreased ADG (*P*<0.05) and ADFI (*P*<0.001) except for the broilers fed with FOS at the stage of challenging with *E. coli* O_78_ (from the 14^th^ day to the 21^st^ day) and the whole trial period in relative to n-control. Moreover, FOS improved the bursal index compared with p-control (*P*<0.05) and had no difference in relation to the n-control. The data demonstrated that *E. coli* O_78_ cause the broilers lose weight and damaged bursa of fabricius. In addition, the Kaplan–Meier curves show the survival of broilers during the period of challenging with *E. coli* O78, the result demonstrated that *L. plantarum* 15-1 and the mixture of *L. plantarum* 15-1 and FOS have reduced the mortality in related to the p-control group, but failed to be guaranteed to no mortality compared with n-control (Figure 1).

**Table 2.**
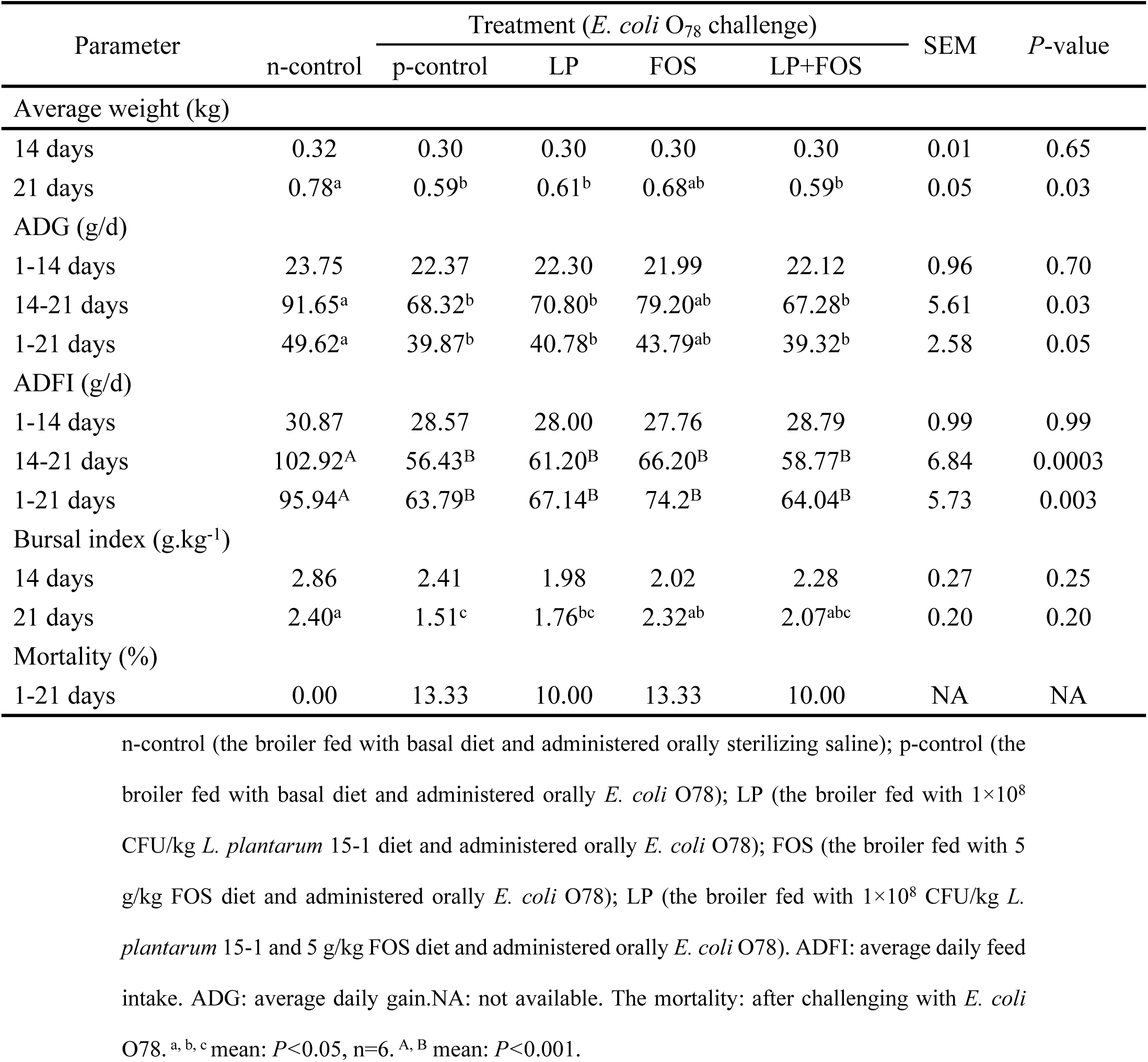
Effect of dietary treatments on performance and mortality of broilers

**Figure 1.**
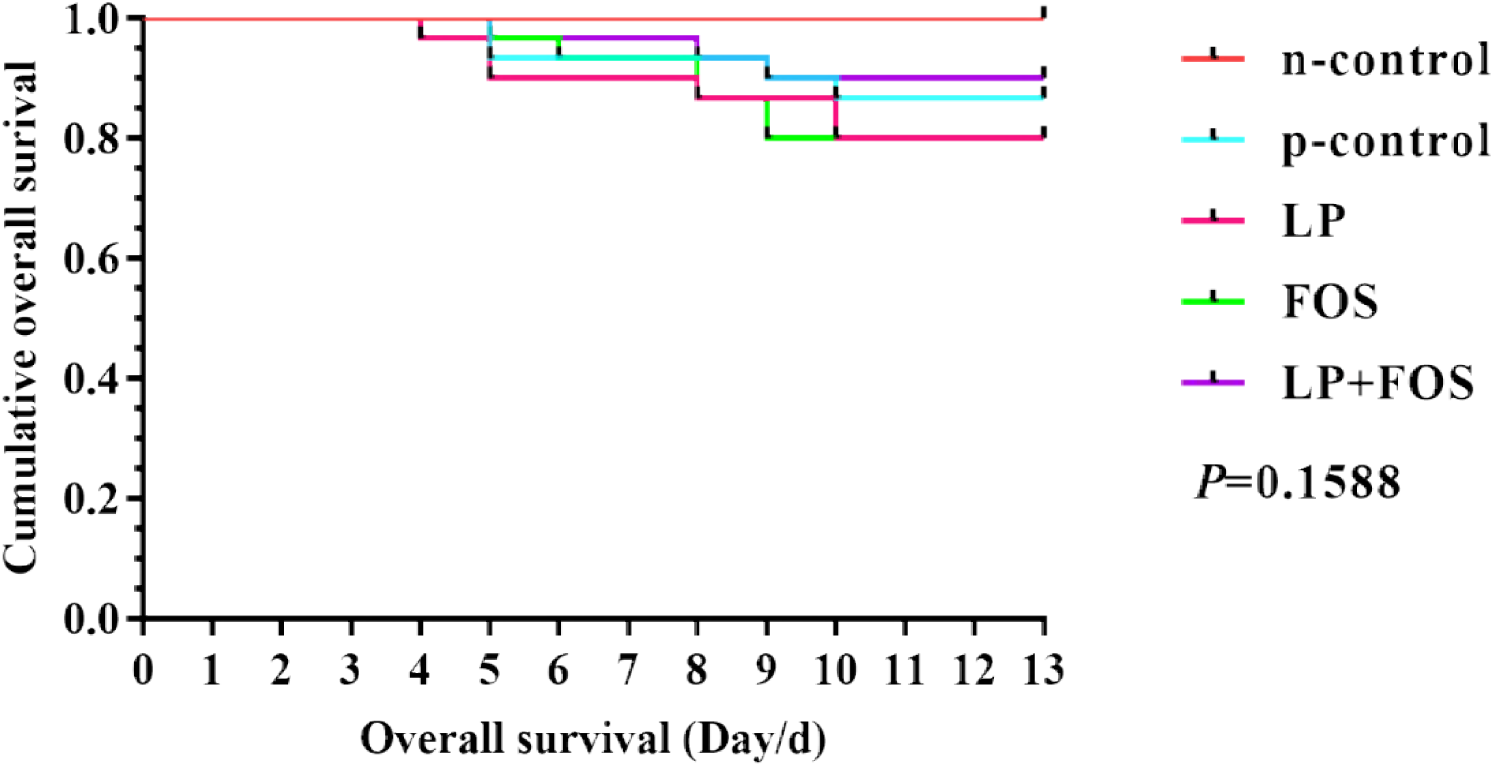
Kaplan–Meier curves displaying survival of broiler began with *E. coli* O78 challenge for 13d. The broiler were fed their respective diets for 7 days, then oral administered *E. coli* O78, while n-control received sterilizing saline (n=30, *P*>0.05). n-control (the broiler fed with basal diet and administered orally sterilizing saline); p-control (the broiler fed with basal diet and administered orally *E. coli* O78); LP (the broiler fed with 1×10^8^ CFU/kg *L. plantarum* 15-1 diet and administered orally *E. coli* O78); FOS (the broiler fed with 5 g/kg FOS diet and administered orally *E. coli* O78); LP (the broiler fed with 1×10^8^ CFU/kg *L. plantarum* 15-1 and 5 g/kg FOS diet and administered orally *E. coli* O78).

### Jejunal Morphology

*L. plantarum* 15-1 or FOS decreased (*P*<0.05) crypt depth in comparison with n-control and p-control on d 14. On d 21, *L. plantarum* 15-1 or FOS, *L. plantarum* 15-1 and FOS lowered crypt depth in comparison with p-control (*P*<0.05) but had no difference relative to n-control (Figure 2). There was no other significance difference.

**Figure 2.**
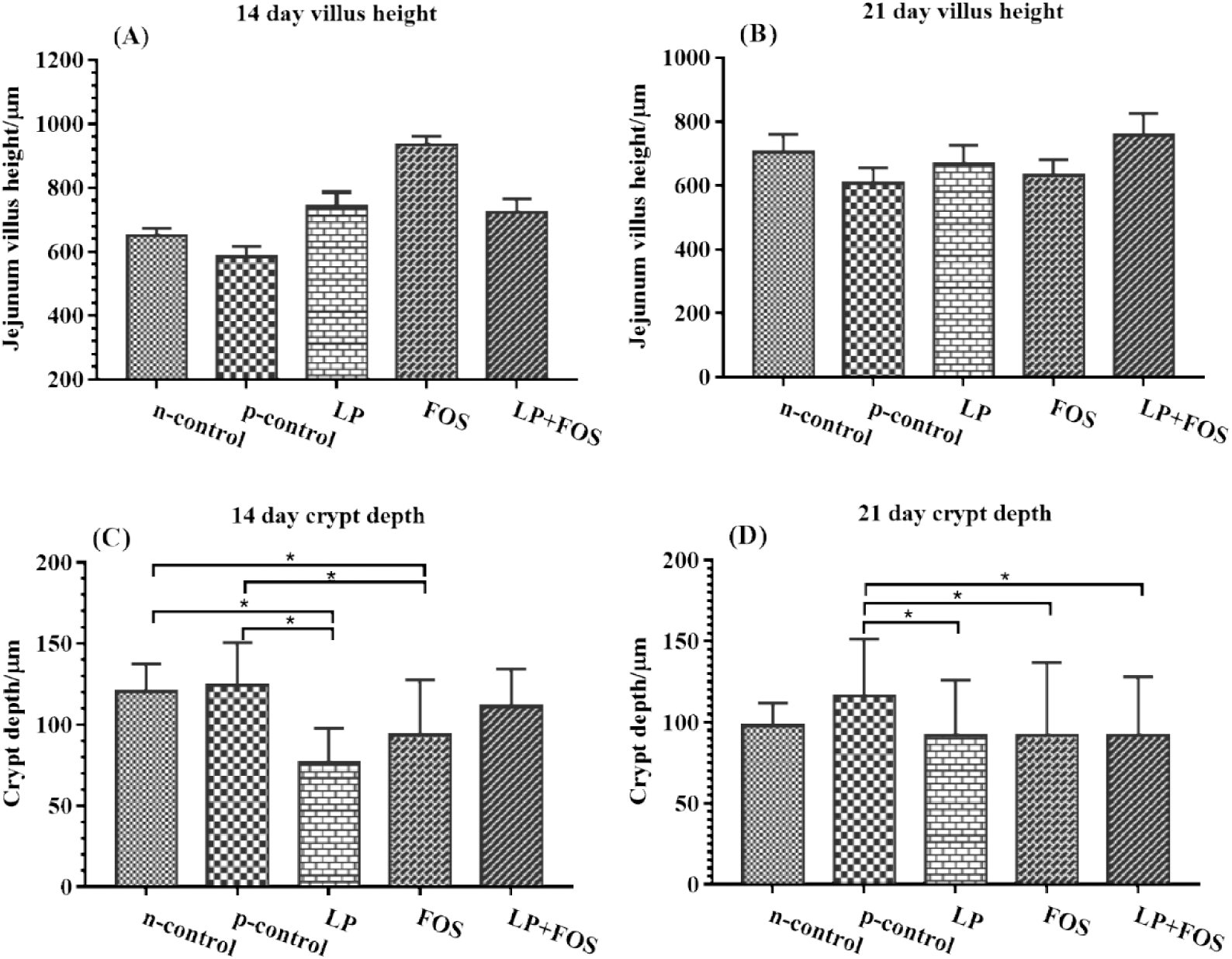
Effect of dietary treatments on intestinal morphology of jejunum of broilers after *E. coli* O78 challenge. n-control (the broiler fed with basal diet and administered orally sterilizing saline); p-control (the broiler fed with basal diet and administered orally *E. coli* O78); LP (the broiler fed with 1×10^8^ CFU/kg *L. plantarum* 15-1 diet and administered orally *E. coli* O78); FOS (the broiler fed with 5 g/kg FOS diet and administered orally *E. coli* O78); LP (the broiler fed with 1×10^8^ CFU/kg *L. plantarum* 15-1 and 5 g/kg FOS diet and administered orally *E. coli* O78). * indicates *P* < 0.05.

### Immune Responses

The impact of *L. plantarum* 15-1 and FOS on the sero-immunity level showed in Figure 3, *L. plantarum* 15-1 reduced DAO (*P*<0.001) compared with p-control, other showed no difference in day 14. *L. plantarum* 15-1 and FOS increased the level of Ig A in relative to p-control (*P*<0.001), while *L. plantarum* 15-1, *L. plantarum* 15-1 and FOS increased the level of Ig G in comparison with p-control, and reduced DAO compared with two control groups in day 21 (*P*<0.001).

**Figure 3.**
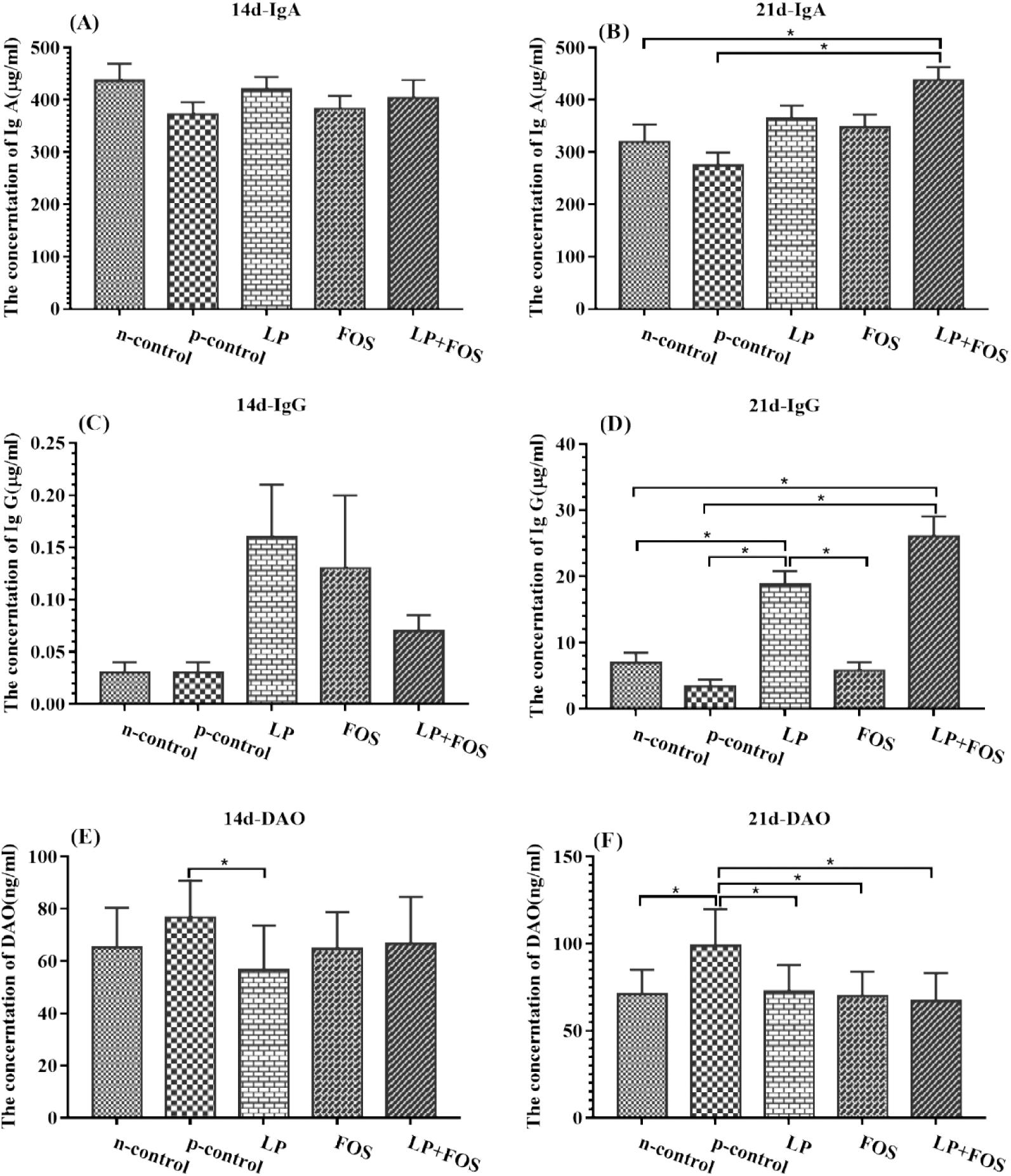
Effect of dietary treatments on concentrations of IgA, IgG and DAO in the serum of broilers. n-control (the broiler fed with basal diet and administered orally sterilizing saline); p-control (the broiler fed with basal diet and administered orally *E. coli* O78); LP (the broiler fed with 1×10^8^ CFU/kg *L. plantarum* 15-1 diet and administered orally *E. coli* O78); FOS (the broiler fed with 5 g/kg FOS diet and administered orally *E. coli* O78); LP (the broiler fed with 1×10^8^ CFU/kg *L. plantarum* 15-1 and 5 g/kg FOS diet and administered orally *E. coli* O78). * indicates *P* < 0.05.

### Cecal short-chain fatty acid

The caecum content was analyzed by gas chromatography after unfroze at 4 °C. *L. plantarum* 15-1 enhanced the level of acetic acid and total SCFA relative to p-control (*P*< 0.05), while increased the level of butyric acid compared with n-control in day 14. FOS increased the level of acetic acid and total SCFA compared with two control groups, while increased butyric acid in comparation to n-control in day 14 (*P*<0.05), moreover FOS increased the level of valeric acid compared with p-control (*P*<0.001). In addition, there show no difference in the broiler fed with *L. plantarum* 15-1 and FOS (Figure 4).

**Figure 4.**
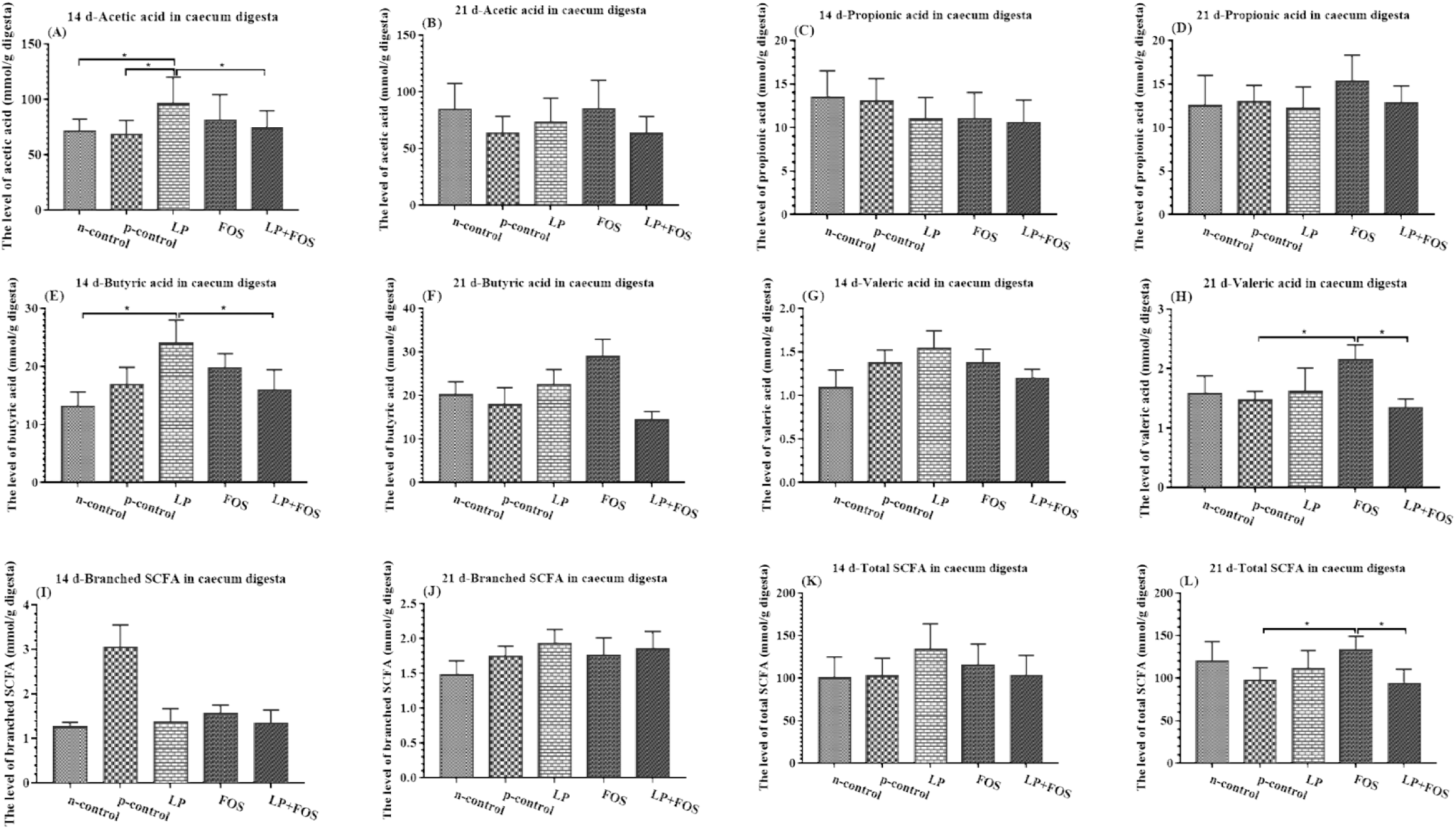
Effect of dietary treatments on short chain fatty acids of broilers at 14 and 21 days of age. n-control (the broiler fed with basal diet and administered orally sterilizing saline); p-control (the broiler fed with basal diet and administered orally *E. coli* O78); LP (the broiler fed with 1×10^8^ CFU/kg *L. plantarum* 15-1 diet and administered orally *E. coli* O78); FOS (the broiler fed with 5 g/kg FOS diet and administered orally *E. coli* O78); LP (the broiler fed with 1×10^8^ CFU/kg *L. plantarum* 15-1 and 5 g/kg FOS diet and administered orally *E. coli* O78). * indicates *P* < 0.05.

## DISCUSSION

It is great significance for improving poultry health to reduce the immune and intestinal damage caused by pathogenic *Escherichia coli*. FOS is conducive to the growth of animals and resistance to pathogenic bacteria. FOS is a preferable substrate to improve the growth of *Bifidobacteria* and promote it combine with the host mucous membrane, which hinders the binding of pathogenic bacteria to the intestinal tract of the host [22]. The result of this study has demonstrated that *L. plantarum 15-1* and FOS could ameliorate the intestinal damage induced by *E. coil* O78 and increased the immune responses, such as improving the level of IgA and IgG and reducing the concentration of DAO in serum. Moreover, *L. Plantarum 15-1* and FOS increased the level of SCFA in cecal contents to resist the invasion of pathogenic bacteria. Those result revealed that *L. Plantarum* 15-1 and FOS could maintain the health status of the broilers.

*E. coli* is associated with the deterioration of animal health, including weight loss, diarrhea, mortality, and necrotizing enteritis [23]. Gaëlle Porcheron’s [24] research shows that FOS may be capable of regulating an extra-intestinal pathogenic *Escherichia coli* strain, and this property is associated with a gene cluster called the *fos* locus, plays a major intestinal colonization, which brings good results that the probiotic bacteria of the microbiota can metabolize in the intestine and decrease the population of pathogenic bacteria. However, no significant differences were found in growth performance. The same discoveries have also been found in other studies, supplementation 0.5% didn’t effect on the growth performance of broiler. On the contrary, there is a proof of study that FOS enhanced FCR but lowered the feed intake and daily live weight gain without challenging with *Escherichia coli* [25]. G.-B. Kim et al. reported that 0.25% FOS as the supplement improved growth performance of the broiler at 28 d [16]. Furthermore, Xu et al. [26] found that FOS increased body weight and feed conversion ratio, and the effect of FOS was much better in 3-week-old than 1-week-old, this result may cause that it takes time to utilize for the beneficial groups and grow into the dominant microflora to keep the balance of intestinal microbial environment and then improve the body growth of broilers with *Escherichia coli* challenge. The results of the present study were in agreement with Xu et al. Many bacteria are used as probiotic, involving *Lactobacillus*, *Leuconostoc*, *Pediococcus*, *Bifidobacterium*, and *Enterococcus*, which produce a promoted action for the growth of animals [27, 28]. Q. Peng et al. had found that *Lactobacillus Plantarum* B1 higher ADG during the finisher period. A Saffar et al. [29] studies probiotics to reduce broilers ascites in high altitude areas, the results found that probiotics reduce the ascites mortality has a role in promoting, but not to compensate for the growth of a strong impact. This may indicate that *Lactobacillus Plantarum* and FOS improve the level of broilers performance. This study has showed that *L. Plantarum 15-1* and FOS deduced the morality of mortality after challenging with *E. coil* O78. Moreover, *L. Plantarum 15-1* and FOS treatment decreased the crypt depth on day 14 and 21. But there no difference on weight compared the p-control group.

It is necessary for poultry to keep the intestinal tract healthy to absorb nutrients and act as a barrier against pathogenic bacteria. Awad W A et al. [30] indicated that *Lactobacilli* has a positive effect on the gastrointestinal tract on this account that it increases feed consumption and absorption of nutrient in intestinal architecture. The intestine is the important site of nutrient absorption, and its efficiency is concerned with its surface area due to an increasing villus and mucosa thickness [31]. A Saffar et al. [32] studies probiotics to reduce broilers ascites in high altitude areas, the results found that probiotics reduce the ascites mortality has a role in promoting, but not to compensate for the growth of a strong impact. Challenge of *E. coli K88* caused to the atrophy of the villi and the destruction of intestinal morphological [33], Pan L et al. [33]indicated that the probiotic has the capacity to protect intestinal barrier function from enterotoxigenic *Escherichia coli* K88 and also showed the probiotic could be a potential candidates to replace the antibiotics for improving the diarrhea. Awad et al. [34]the broiler diets supplied with Lactobacillus increased villus height and villus height: crypt depth ratio, and Song J et la. [35]reported that *Lactobacillus Plantarum* as mixed probiotics with *Bacillus licheniformis* and *Bacillus subtilis* increased the jejunal villus height and decreased small intestinal coliforms. The present study have indicated that *L. plantarum* 15-1 added to based diet contributes to the tissue morphology of jejunum.

DAO can reflect the structure and function of the mucosa integrity and small intestinal mucosa. under normal circumstances, the level of DAO in serum are very low, but the concentration of DAO increase after intestinal mucosa is damaged [36]. Several studies have shown that the addition of probiotic and prebiotic improve the host immunity via increasing the concentration of IgA, Kim C H et al. [37] found that the diet supplied with FOS could increase on laying hens. On the contrary, Kim G B et al. [16] reported that the concentration of plasma IgA and IgG in broiler plasma had no different between the broilers fed diet with the supplement of FOS and other treatment groups. Maragkoudakis P A et al. [38]demonstrated that no differences were found in the IgA, IgM and IgG levels in goat plasma added with FOS. Also Gürbüz E et al. [39] demonstrated there is no difference in the concentration of IgA, IgM and IgG in horses. In this study, the result showed that *L. plantarum* 15-1, FOS and their mixed treatment increased the serum levels of IgA and IgG in d 21 and decreased DAO.

Short-chain fatty acids are the pivotal metabolite of microorganism during fermentation of indigestible carbohydrates in the large intestine and used by the colonic mucosa [12, 40]. Lactic acid bacteria enhance the physical barrier of the host via increasing the SCFA, and high level of SCFA in the environment is not conducive to the growth and reproduction of pathogenic bacteria. SCFA is the final products of microbial fermentation that is absorbed in the colonic mucosa [41]. SCFAs is an ideal biorenewable chemicals to inhibit E. coli that maybe biorenewabel chemical damage the cell membrane of the pathogen [42]. Application of prebiotics enhanced the level of SCFAs, thus improving intestinal acidity, and that is conducive to lower pathogens in the gut of chicken. [43]. Butyrate is a direct source of energy for the colonic epithelium, equipped with anti-inflammatory properties and can enhance the colonic defence barrier [44]. Peng Q et al. [45] reported that *L. plantarum* B1 increased the SCFA in the cecum. In the present study, the total SCFA was increased by *L. plantarum* 15-1, FOS and their mixed, especially acetic acid and butyric acid in d 14 and valeric acid in d 21. This may be one of the reasons that probiotics reduce mortality. Thoughtfully, there are study showed that synbiotics couldn’t show the double effect of two or more substances regarding growth performance, intestinal microbial, and the level of SCFA [46]. This discrepancy maybe contributed to the different properties of prebiotics and probiotics.

In conclusion, supplementation with FOS and *L. plantarum* 15-1 improved intestinal morphology, enhanced immune responses and increased the concentration of SCFA in cecum challenged with *E. coli* O_78_. Moreover, *L. plantarum* 15-1 and FOS had no effect on growth performance but decreased the mortality of the broilers challenged *E. coli* O_78_. That’s all demonstrated FOS and *L. plantarum* 15-1 may could ameliorate the negative effect of *E. coli* O_78_.

## Ethics statement

All animal procedures were approved by the Animal Ethics Committee Guidelines of Academy of State Administration of Grain, Beijing, China (20170052), and performed according to the guidelines recommended in the Guide for the Laboratory Animal Ethical Commission of State Administration of Grain.

## Author contribution

Experimental design: Sujuan Ding, Yongwei Wang;

Animal feeding: Sujuan Ding;

Detection of indexes: Sujuan Ding, Wenxin Yan;

Analysis of data: Wenxin Yan, Hongmei Jiang;

Writing – original draft: Sujuan Ding;

Writing – review & editing: Jun Fang, Aike Li;

Provision of funds: Jun Fang, Aike Li.

